# The widespread nature of Pack-TYPE transposons reveals their importance for plant genome evolution

**DOI:** 10.1101/2021.06.18.448592

**Authors:** Jack S. Gisby, Marco Catoni

**Author notes:** **Contact details:** Marco Catoni, Jack S. Gisby. Department of Immunology and Inflammation, Imperial College London, London, United Kingdom, W12 0NN (Jack S. Gisby).

## Abstract

Pack-TYPE transposable elements (TEs) are a group of non-autonomous DNA transposons found in plants. These elements can efficiently capture and shuffle coding DNA across the host genome, accelerating the evolution of genes. Despite their relevance for plant genome plasticity, the detection and study of Pack-TYPE TEs are challenging due to the high similarity these elements have with genes. Here, we produced an automated annotation procedure designed to study Pack-TYPE elements and used it to successfully annotate and analyse more than 4000 new Pack-TYPE TEs in the rice and maize genomes. Our analysis indicates that Pack-TYPE TEs are an abundant and heterogeneous group of elements. We found that these elements are associated with all main superfamilies of Class II DNA transposons in plants and likely share a similar mechanism to capture new chromosomal DNA sequences. Furthermore, we report examples of the direct contribution of these TEs to coding genes, suggesting a generalised and extensive role of Pack-TYPE TEs in plant genome evolution.

## Introduction

Class II DNA transposable elements (TEs) are genomic loci able to move their sequence and relocate it to a new chromosomal location. This mobilisation process is mediated by a transposase typically encoded in the TE sequence, able to specifically recognise Terminal Inverted Repeat DNA sequences (TIRs) located at each end of the TE (1–3). TEs that lack a functional transposase gene are defined as non-autonomous, and they can transpose if a transposase is provided *in trans* by a related autonomous element, as long as they maintain functional TIRs (4). Due to reduced constraints on their DNA sequence, non-autonomous TEs can often become more abundant than their corresponding autonomous counterpart (5, 6).

Pack-TYPE elements are a kind of Class II non-autonomous TE discovered in plants that contain DNA fragments captured from coding genes between their TIR sequences (7, 8). These elements can acquire and rearrange host genes during mobilisation, a process called transduplication. Pack-TEs can potentially affect gene expression and increase gene diversity (7, 9). In rice (*Oryza sativa*), more than 3000 Pack-TYPE TEs were annotated and classified as Pack-MULEs due to their TIR homology to autonomous elements belonging to the *Mutator*-like (MULE) superfamily (8). However, the presence of coding gene fragments and the high variability in the sequence of these elements represent a challenge for automatic TE annotation tools. Although Pack-MULEs have since been found in other plants (10, 11), specific annotation approaches were required to identify their sequences in the genomes.

Recently, in *Arabidopsis thaliana*, a new family of Pack-TYPE elements has been discovered with TIRs similar to autonomous elements of the CACTA superfamily, which were therefore defined Pack-CACTA (7). In contrast to Pack-MULEs, Pack-CACTAs mobilise in Arabidopsis epigenetic recombinant inbred lines, and their study contributed to clarifying their transposition process and the mechanism used to acquire new coding DNA (7). Although it was unclear how common Pack-CACTAs were in plant genomes, their discovery in Arabidopsis demonstrated that MULE is not the only superfamily with Pack-TYPE TEs; this raised the question of whether other superfamilies of TEs with TIRs could potentially support the transposition of compatible Pack-TYPE elements (12).

Here, we use common features of Pack-TYPE transposons to automatically annotate these elements in plant genomes, providing a specific and reliable tool for their annotation and clustering. In addition, we show that Pack-TYPE TEs are not limited to the MULE and CACTA superfamilies, and we found examples of exon shuffling events mediated by Pack-TYPE TEs belonging to all main superfamilies of plant DNA transposons with TIRs.

## Methods

### Reference genomes

The *Arabidopsis thaliana* TAIR reference genome (GCF_000001735.4), the *Oryza sativa* IRGSP-1.0 reference genome (GCF_001433935.1) and the *Zea mays* B73 reference genome (GCF_000005005.2) were downloaded from NCBI. In addition, general TE annotations for the *A. thaliana, Zea mays* and *Oryza sativa* genomes were obtained respectively from TAIR10 (https://www.arabidopsis.org/), the maize B73 annotation (https://mcstitzer.github.io/maize_TEs), and the rice IGRSP1 annotation project (https://rapdb.dna.affrc.go.jp).

### Automatic detection of Pack-TYPE transposons

To automatically detect Pack-TYPE TEs, we implemented an algorithm in three steps. In the first step, we used conserved TIR reference sequences of 8 to 13 bp at the terminal ends of well characterised autonomous TEs to survey the reference genome and find matches (**Table 1**). Base pair mismatches and indels are allowed by calculating Levenshtein distance to find optimal local matches (13). We report the reference TIRs used and the number of allowed mismatches for each analysis in **Table 1**. The genomic fragments delimited by TIR pairs found close within the genome (in the interval of 300-15,000 bp) and in inverted orientation were selected as tentative TEs. This list was then filtered based on the presence of a TSD and its similarity at the 5’ and 3’ of the putative TE, again calculated using Levenshtein sequence distance (**Table 1**). Duplicated or overlapping TIR reference sequences can be defined due to the use of multiple TIR reference queries (**Table 1**); these were identified using the GenomicRanges (14) package and removed to prevent overestimation of TE abundances.

**Table 1.**
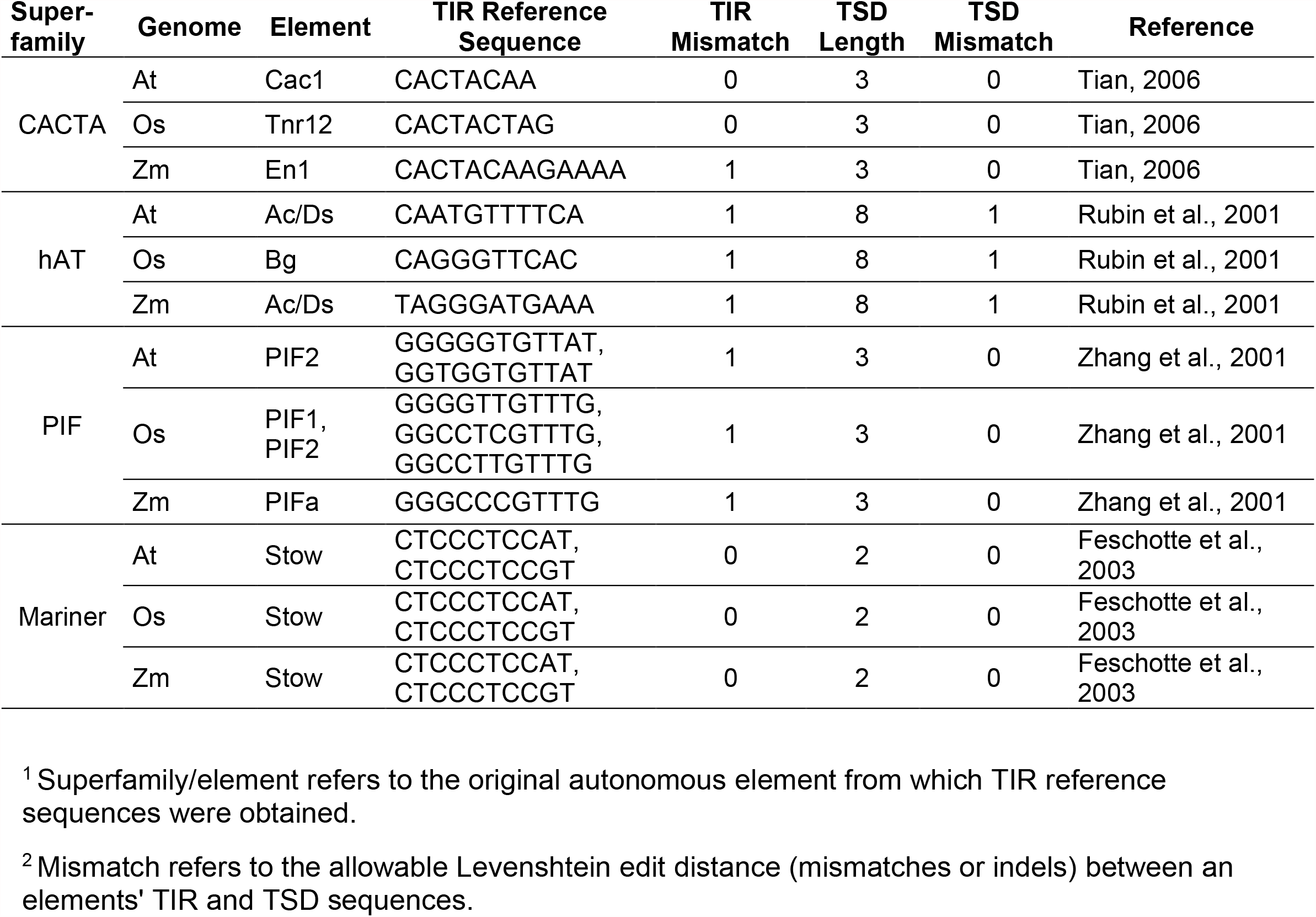
TIR sequences and parameters used to identify Pack-TYPE elements. (*At* = *Arabidopsis thaliana*; *Os* = *Oryza sativa*; *Zm* = *Zea mays*).

Subsequently, in the second step, we clustered (centroid based) tentative TEs using VSEARCH (15) to group TEs belonging to the same family. VSEARCH (default parameters) clustered DNA sequences together where the edit distance was greater than 60%. To ensure quality alignments and stringent selection of putative TEs, we removed sequences with >10% wildcard nucleotides (N). To control for false positives and generate reliable lists of potential TEs, we removed singleton clusters under the assumption that real TEs are repeated in the genome. We also assumed that elements inside each cluster are evolutionarily linked, excluding the possibility of independent capture events of the same chromosomal sequence in different TEs (that should be rare). Therefore, in this work, we consider the cluster to be the closest approximation of a TE family. Considering that the orientation of non-autonomous TEs cannot be defined easily, we arbitrarily considered the largest element of each cluster as forward orientated, and we determined the relative direction of the remaining TEs belonging to the same cluster by alignment.

The final list of putative TEs could include autonomous and non-autonomous TEs, and Pack-TYPE or not Pack-TYPE elements. Therefore, we applied a third step to automatically classify the putative TEs and identify potential Pack-TYPE TEs, according to the categories previously applied to Pack-MULEs (10). Specifically, we used the CDS from the relevant reference genome (*Arabidopsis thaliana, Zea mays* or *Oryza sativa)* and the BLAST tool (16, 17) to identify elements matching the sequence of transposase proteins. Any putative TE with a BLAST match (evalue < 1e-5, length > 250) to a transposase was automatically classified as an “ autonomous TE” and was considered to be autonomous or derived from an autonomous element. We categorised the remaining elements as Pack-TYPE or non-Pack-TYPE based respectively on the presence or absence of BLAST hits (evalue

< 1e-5, length > 50) to a valid CDS entry. In this step BLAST was run on CDS databases with the options -max_target_seqs 500, -task blastn-short and -word_size 7 (7). For each annotated TE of a specific superfamily and genome, we assigned a numerical ID in the order of discovery. To generate a unique element identifier, we concatenate the genome, the superfamily of the TIR sequence used for detection and the assigned ID (e.g. “ At-CACTA-5” or “ Os-Mariner-12”). We implemented the entire procedure in an R package called *packFinder* (18), which is publicly available as part of the Bioconductor project (19). The results presented here were generated using *packFinder* v1.2.0.

### Analysis of Neighbouring Elements

For each cluster of TEs, we calculated the proportion of cluster elements within 100kb of another member of the cluster and compared this to the proportion of all non-member TEs within 100kb of a member of the cluster. We then tested the two proportions using a one-sided Chi-squared test for each TE superfamily. The directionality of annotated TEs was defined by their orientation relative to the largest element of their respective clusters.

### TIR Clustering

We obtained the first 80 bp of all identified Pack-TYPE TE sequences, representing the forward TIRs, and used these to generate a distance matrix, using alignment-free kmer counting and a kmer size of 5 (default) (20). We then applied quantile-based colour breaks to visualise the matrix as a heatmap, and complete-linkage hierarchical clustering to order the columns and rows of the distance matrix.

### Mappability

Mappability can estimate the repetitiveness of a TE in a genome (21). Here, we computed mappability for both the rice and maize genomes with the GemMap tool (22), using a size of 20 and allowing a maximum of one mismatch. We converted the GemMap output to bigwig format in R with the package rtracklayer (23). We then used deepTools (24) to plot the distribution of mappability averaged for each TE group, using a bin size of 20 nt. We obtained the general annotation of maize CACTA, PIF and hAT TEs using the B73 maize repeats annotation filtered for elements classified as “ DTC”, “ DTH”, and “ DTA”, respectively. The annotation of rice Mariner TEs was obtained by extracting “ Stowaway” elements from IRGSP1 repeats annotation (25).

### Synteny

We selected Pack-TEs overlapping gene coding regions using the R GenomicRanges package (14). Then, syntenic genomic regions were identified using the EnseblPlants Comparative Genomics tools (https://plants.ensembl.org/index.html) in phylogenetically related species. The DNA sequence of the relevant loci were downloaded and aligned with Geneious (https://www.geneious.com) to generate alignment scores.

## Results

### Automatic annotation of Pack-TYPE TEs

In previous analyses, Pack-MULE and Pack-CACTA elements inserted respectively in the *Oryza sativa* and *Arabidopsis thaliana* reference genomes were identified using a combination of BLAST and manual annotation (7, 8). In order to optimise and automate the annotation of Pack-TYPE transposons for metagenomics studies, we standardised the detection procedure in an algorithm implemented in the R package *packFinder* (18). The algorithm takes as input TIR sequences and categorises the identified putative TEs according to the functional annotation previously applied to Pack-MULEs (10), which consists of three groups: i) autonomous TEs (with a valid blast hit to a transposase); ii) Pack-TYPE TEs (with a valid BLAST hit to coding genes); iii) non-Pack-TYPE TEs (remaining elements without significant blast hits to DNA transposases or coding genes).

Using as input the TIRs of the *Arabidopsis thaliana* autonomous CACTA elements (**Table 1**) (26), we annotated 54 CACTA-like elements grouped in 12 clusters in the *Arabidopsis thaliana* reference genome (**Table S1**). This list includes the three previously identified Pack-CACTA families with intact TSD sequences (7) found in our analysis as clusters #31, #35 and #40, and two additional clusters (#28 and #39) of non-Pack TEs not previously annotated. The algorithm also found 7 clusters (#2, #6, #9, #10, #14, #22 and #23) of potentially autonomous CACTA TE families. All autonomous TEs identified were already annotated in the TAIR10 database as members of the CACTA superfamily (ATENSPM in Arabidopsis) (**Figure 1A**). However, some Pack-CACTA and non-Pack-TYPE annotations in TAIR10 were inaccurate (i.e. with only a portion of their sequence recognised as CACTA element) (**Figure 1B**), while others were misannotated as a protein coding gene (**Figure 1C**) or as TE repeats not belonging to the CACTA family (but mostly as ATREP and HELITRON families) (**Figure 1D**). This indicates that TAIR10 is not reliable for the annotation of Pack-CACTAs or other CACTA-derived non-autonomous elements. Instead, our automatic algorithm, using only TIR sequences and the Arabidopsis genome as input, correctly annotated intact Pack-CACTA TEs previously manually identified in Arabidopsis, detecting also additional non-Pack-TYPE elements (**Table S1**).

**Figure 1.**
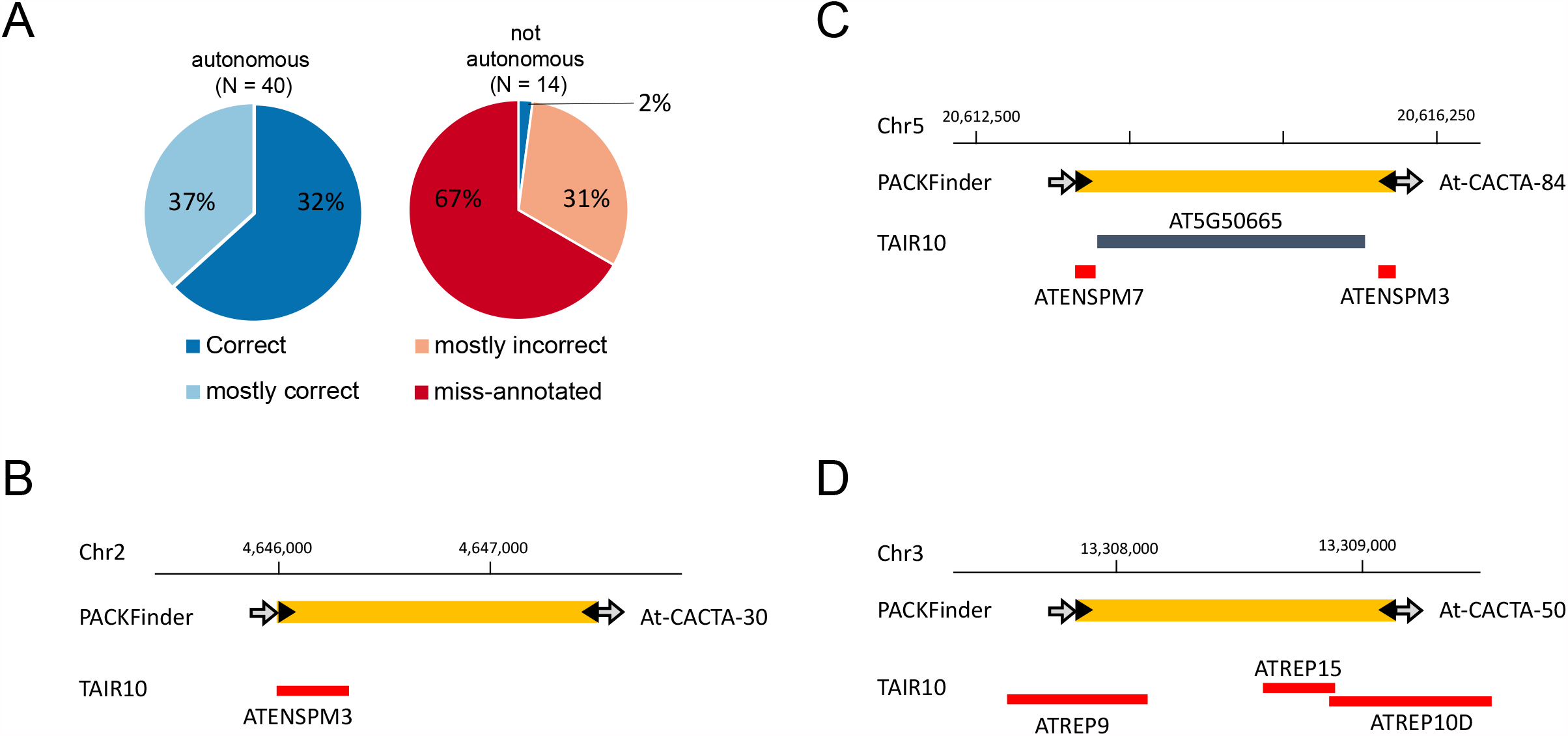
The *packFinder* automatic annotation pipeline identifies Pack-CACTA TEs in Arabidopsis. **A** Pie charts displaying the quality of TAIR10 annotation of autonomous and not autonomous (Pack-TYPE and not Pack) TEs identified by the *packFinder* automatic pipeline. The TAIR annotation have been classified as: i) correct, if the TE is annotated identically by TAIR10 and *packFinder*; ii) mostly correct, if in TAIR10 the TE is not uniquely identified but > 50% of its sequence is annotated as belonging to CACTA (ATENSPM) superfamily; iii) mostly incorrect, if in TAIR10 less than 50% of TE sequence is annotated as belonging to CACTA (ATENSPM) superfamily; or iv) misannotated, if in TAIR10 the TE is not annotated or incorrectly annotated as a coding gene or as belonging to an unrelated repeat. **B, C** Examples of incorrect TAIR10 annotation or **D** misannotation of Pack and non-Pack TEs. TAIR10 annotation as genes (blue box) and TEs (red boxes) is displayed and the corresponding gene name or TE family is reported. *packFinder* annotations (yellow boxes) are displayed with TIRs (black arrow) and TSDs (grey arrows).

### Pack-TYPEs are prevalent across Class II TE superfamilies

Considering that *Arabidopsis thaliana* has a relatively small genome with a low proportion of TEs (27), we applied our automatic annotation workflow to investigate the presence of Pack-TEs in other plants. Moreover, except for the MULE superfamily (8), other TIR TEs (6) have not been comprehensively analysed to identify Pack-TYPE elements. Therefore, to uncover the widespread nature of Pack-TYPE TEs, we investigated whether other DNA TE superfamilies (i.e. hAT, Harbinger-PIF, Mariner) could generate Pack-TYPE elements in rice (*Oryza sativa)* and maize (*Zea mays)*.

To *de novo* annotate CACTA elements, we used the conserved TIR sequences of rice CACTA and maize *En* TEs (26) in our automatic procedure to survey respectively the *Oryza sativa* and *Zea mays* reference genomes. To annotate hAT TEs, we used the core TIR sequences of the well characterised autonomous *Bg* and *Ac* (26, 28) elements for maize and rice, respectively (**Table 1**). Similarly, for the Harbinger-PIF and Mariner superfamilies, we used the core TIR sequences from autonomous *PIF* and *Stow* (Mariner) elements (29, 30) for both *Zea mays* and *Oryza sativa* (**Table 1**). We also applied our algorithm to annotate TEs in the *Arabidopsis thaliana* genome, using TIRs of elements previously described in this species (**Table 1**). However, since we did not detect any new superfamilies of Pack-TE, we excluded this plant from subsequent analyses.

Contrary to Arabidopsis, we successfully annotated TEs for all tested superfamilies in maize and rice (**Table S2**), which appear to be distributed evenly in the genomes (**Figure S1**). In maize, hAT and Harbinger-PIF elements were the most numerous (1,641 and 1,036, respectively), while we found a moderate number of CACTA elements (324 TEs) and relatively few Mariner (52 TEs). Conversely, in the rice genome, we found a high proportion of Mariner elements (642 TEs) and a moderate number of Harbinger-PIF elements (225 TEs), while CACTA and hAT TEs were of lower abundance (75 and 59 elements, respectively).

As expected, all TEs annotated in our analysis contain TIRs and a TSD, consistent with the features of the associated superfamily. Specifically, in both species, all the identified hAT TEs contained a TSD of 8 nt while the CACTA elements had TSDs of 5 nt, as described previously for these groups (26, 28). In addition, the TSDs of virtually all detected Mariner elements were “ TA” in both *Oryza sativa* (99.1%) and *Zea mays* (98.1%), as previously described for this class of elements (29). Similarly, the majority of identified PIF elements had “ TTA” or “ TAA” TSDs (30) in both *Oryza sativa* (52.4%) and *Zea mays* (81.2%). To further confirm the correct annotation of Pack-TYPE TEs, we aligned and clustered the first 80 bp of the forward TIR sequence and observed that, as expected, elements are grouped almost exclusively based on the superfamily to which they belong (**Figure S2**). These observations collectively indicate that our pipeline successfully annotated TEs from different families of DNA transposons.

We then analysed the proportion of annotated Pack-TYPE elements in each superfamily and observed that their number varied depending on the superfamily and plant genome considered (**Figure 2A**). Specifically, we observed that Pack-TYPE elements belonging to the CACTA, hAT and PIF superfamilies were more abundant in maize, while Pack-Mariner accumulated more in the rice genome. In addition, the proportion of Pack-TYPE elements we discovered was largely independent of the number of autonomous elements (or non-Pack TEs) found in the same superfamily (**Figure 2A**).

**Figure 2.**
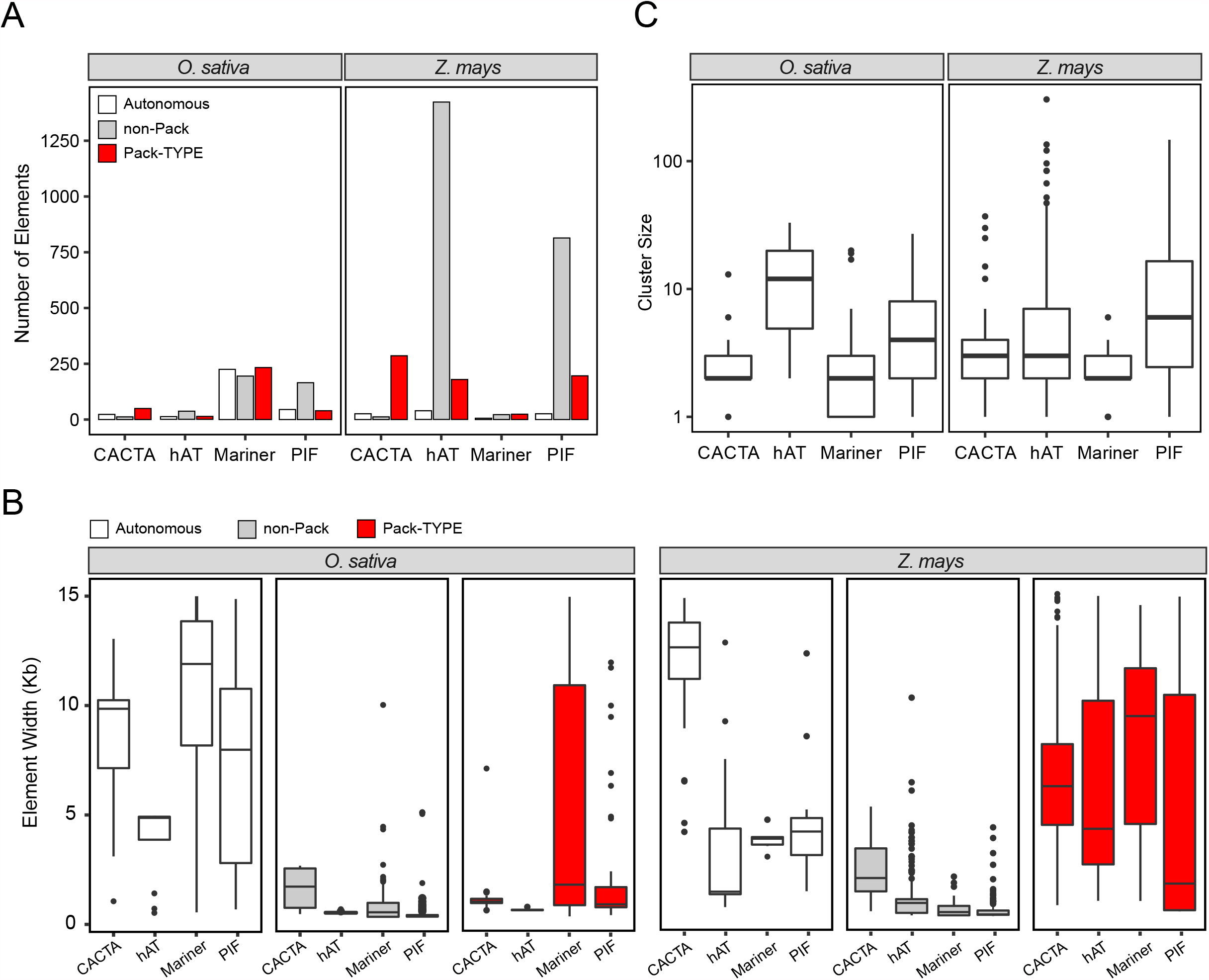
The DNA sequence properties of Pack-TYPE TEs in *Oryza sativa* and *Zea mays*. **A** Bar plot displaying the total number of elements annotated by *packFinder* in each of the TIR superfamilies tested in both rice (*O. sativa*) and maize (*Z. mays*) genomes. Colours represent TE functional designation, assigned automatically using BLAST. **B** Box plots displaying the distribution of TE widths in each category, defined after the application of the *packFinder* BLAST step in both the rice (*O. sativa*) and maize (*Z. mays*) genomes. **C** Boxplots for the distribution of annotated cluster sizes in both rice (*O. sativa*) and maize (*Z. mays*) genomes.

We also observed relatively high variability in the widths of the annotated Pack-TEs, with maize Pack-TYPE elements having a greater median width than rice elements for all annotated superfamilies (**Figure 2B**). In contrast, all TEs classified as non-Pack tended to be smaller than TEs classified as Pack-TYPE or as autonomous in both plant genomes (**Figure 2B**). The cluster size distribution of the annotated TEs (**Figure 2C**) appears to be consistent across the *Oryza sativa* and *Zea mays* genomes, with the largest clusters (containing more than one hundred elements in maize) belonging to the hAT and PIF superfamilies (**Table S2**). By contrast, most CACTA and Mariner Pack-TYPE elements are low copy number in both species (**Figure 2C**).

Similar to rice Pack-MULEs (8) and Arabidopsis Pack-CACTA (7), we also observed examples of Pack-TYPE elements containing genic material from multiple chromosomal loci (**Table S3**). In general, the maize genome appears to contain a higher proportion of elements with coding material captured from multiple loci, while the rice genome contains a greater proportion of elements with coding material derived from single loci; this is consistent with the larger width observed for Pack-TYPE TEs in maize (**Figure 2B**), which can allow for the incorporation of more chromosomal DNA.

### Pack-TYPE TEs acquire new DNA by mobilisation of neighbouring insertions

The frequent excision and insertion of DNA TEs in local chromosomal areas (also called “ local hopping”) has been proposed to facilitate the acquisition of new DNA and the evolution of Pack-CACTA TEs in Arabidopsis (7). We searched for evidence of local hopping occurring during the mobilisation of Pack-TYPE TEs by investigating instances of local insertions in the set of annotated Pack-TYPE clusters.

We defined local insertions as all pairs of annotated TEs belonging to the same cluster and within 100kb of distance from each other. In order to have a sufficient number of pairs to apply statistics, we focussed the analysis only on superfamilies with at least ten local insertions, which included the maize hAT (15 local insertions belonging to 8 clusters) and PIF (13 local insertions belonging to 5 clusters) superfamilies. For both groups, the aggregated proportion of observed local insertions of elements belonging to the same cluster was significantly greater (for hAT-like p= 6.6⨯10^−12^ and for PIF-like p= 2.9⨯10^−11^) compared to the occurrence of local insertions of members of different clusters. We also observed that among all local TE pairs considered, 80% (12 out of 15) for hAT and 92% (12 out of 13) for PIF were in the same orientation on the chromosome, suggesting that these TEs can influence the direction of insertion in its proximity for elements of the same family. Interestingly, local insertions with conserved orientation have been observed in real-time during the mobilisation of Pack-CACTA elements in Arabidopsis (7).

If two Pack-TYPE TEs with compatible TIRs are inserted within a short distance at the same locus, a transposase could recognise the two external TIRs as the start and the end of a mobile element; and the two original TEs can then mobilise as a single element, including the DNA located initially between the two insertions (7) (**Figure 3A**). Therefore, Pack-TYPE TEs should contain newly acquired DNA located internally in phylogenetically related elements, while TIRs and terminal DNA sequences should be more conserved among elements of the same family. We tested this hypothesis by estimating the average uniqueness (calculated with the mappability, see **methods**) of DNA sequences for each superfamily in the genome where most Pack-TYPE TEs were annotated, including Pack-Mariners in the rice genome and Pack-CACTAs, Pack-PIFs and Pack-hATs in the maize genome. We observed that all Pack-TYPEs tested contain less repeated DNA than all TEs annotated in rice and maize belonging to the corresponding superfamily (**Figure S3**), and this effect tended to be more relevant in the internal part of the TEs (**Figure S3**). This result is compatible with a model where more recently acquired coding DNA (assumed to be less repeated) is located more internally and further from TIR sequences.

**Figure 3.**
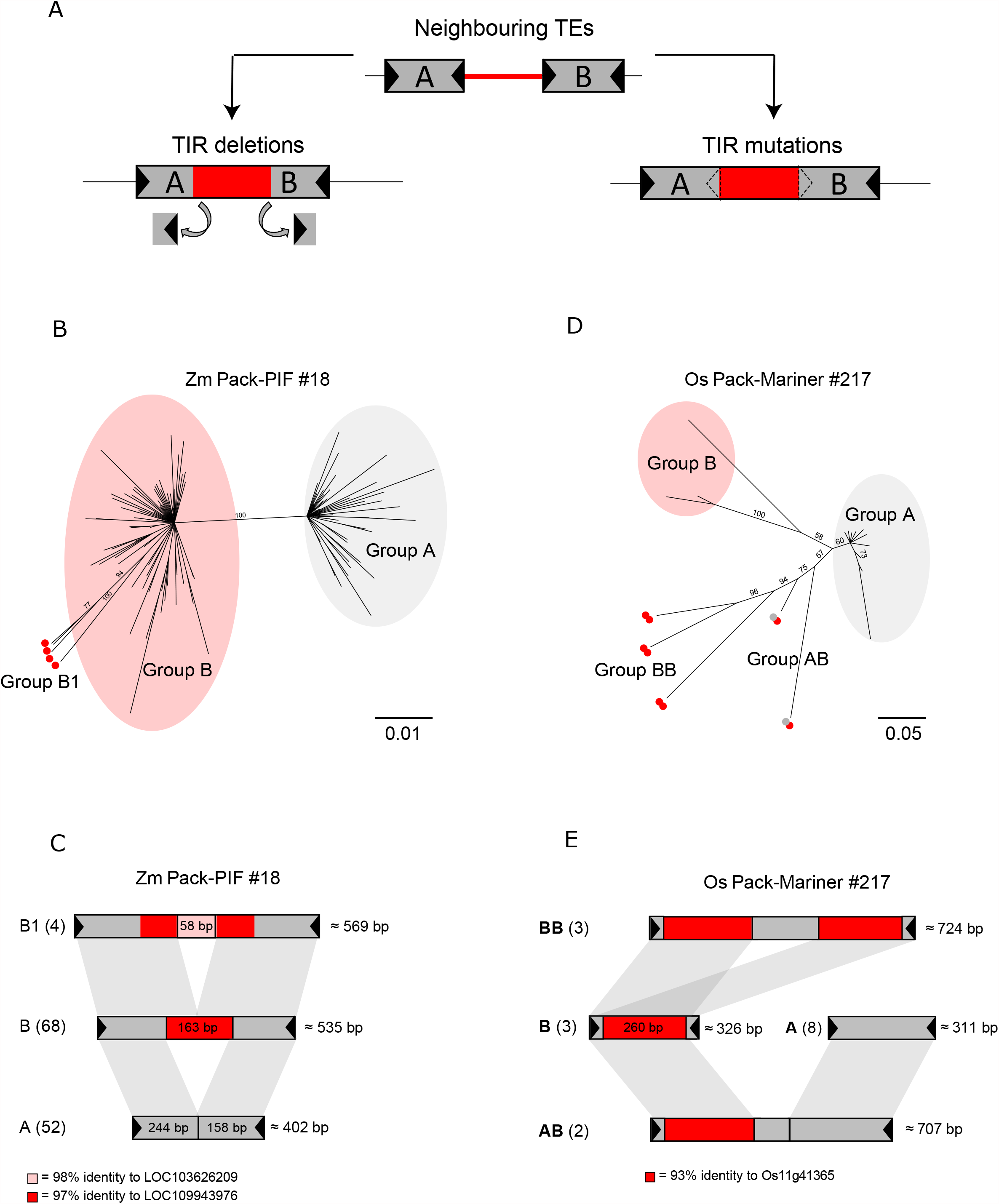
Evidence of Pack-TYPE TE evolution in maize and rice genome. **A** Scheme illustrating the model for the acquisition of chromosomal DNA by Pack-TYPE TEs (Catoni et al., 2019). A pair of neighbouring Pack-TYPE TE insertions (marked as A and B) and the DNA region between them (marked in red) can be recognised as single elements and mobilised by a transposase either by deletion (left) or mutations (right) occurring at the internal TIRs. A black triangle represents TIRs. **B** Phylogenetic tree from alignment of full-length sequences of elements in Zm-Pack-PIF cluster #18. Numbers at each node indicate bootstrap support values of 100 replications. Elements in groups and subgroup are shaded and marked with a red dot, respectively. **C** Structure of the Zm-Pack-PIF TE family #18. Each bar represents a group or subgroup as shown in **B** (number of elements per group indicated in parenthesis). Shaded grey marks regions with sequence homology. The average size of elements in each group is displayed on the right. In red is a 163 bp region with 97% identity to the maize gene LOC109943976, while pink marks a 58 bp region with 98% identity to the gene LOC103626209. The black triangles mark the TIRs (15 nt). **D** Phylogenetic tree from the alignment of full-length sequences of elements in Os-Pack-Mariner cluster #217. Numbers at each node indicate bootstrap support values of 100 replications. Elements in groups A and B are shaded in grey and red, respectively. TEs including two copies of elements in group B are marked with two red dots (Group BB), while elements including a copy of A and a copy of B are marked with a red and a grey dot (Group AB). **E** Structure of Os-Pack-Mariner TE family #217. Each bar represents a group shown in D (with the number of elements per group indicated in parenthesis). Shaded grey marks regions with sequence homology. The average size of elements in each group is displayed on the right. A 260 bp region with 93% identity to the rice gene Os11g41365 is displayed in red. The black triangles mark the TIRs (15 nt).

We further investigated the structure of these Pack-TYPE TEs by alignment at the level of single clusters and found examples of DNA sequence acquisition events that are compatible with the fusion of neighbouring TE insertions. For example, in Zm-Pack-PIF cluster #18 in maize, we classified TEs into two groups based on their sequence similarity (**Figure 3B** and **Table S4**). Group A was composed of 52 non-Pack elements with similar lengths (average size of 402 bp), while Group B constituted 72 slightly longer Pack-TYPE TEs (with an average size of 535 bp). The main difference among elements of the two groups was an internal insertion of 163 bp found in all TEs in group B, with 97% similarity to the gene LOC109943976. In four elements of this group (subgroup B1), an additional insertion of 58 nucleotides was found, with 98% similarity to the gene LOC103626209 (**Figure 3C**).

In a second example, the Os-Pack-Mariner cluster #217 in rice, elements could be separated into four groups (**Figure 3D** and **Table S4**). Group A includes eight non-Pack TEs with an average length of 311 bp, while B contains three Pack-TYPE elements (that include a 280 bp DNA portion with 93% similarity to the rice gene Osg41365) and TIR sequences similar to group A (**Figure 3E**). Interestingly, three elements (Group AB) contain both the sequence of the A and B groups spaced by some additional DNA of unknown origin, while another two elements (group BB) have a similar structure but contain the sequence of group B repeated twice (**Figure 3E**). This suggests that two smaller TEs fused to generate elements of the AB and BB groups.

We observed similar examples of sequence acquisition events for Pack-TYPE TEs belonging to the CACTA and hAT superfamilies (**Figure S4**). Collectively, these observations suggest that Pack-TYPE TEs of all DNA superfamilies can acquire DNA with a similar mechanism based on tandem insertions and re-mobilisation, as previously hypothesised for Pack-CACTA (7).

### Pack-TYPE TEs contribute to plant gene evolution

During mobilisation, Pack-MULE TEs shuffle exons across the genome, contributing to the generation of new transcript variants (8). Genes that evolve in such a way can acquire large portions of coding DNA in a short evolutionary time, and homologs in other species can lack the DNA encoded by the sequence of the Pack-TYPE TEs (11). To test this property in the newly identified Pack-TYPE superfamilies, we retrieved genes with coding sequences that overlap Pack-TYPE TEs (**Table S5**). We then used the Plant Ensembl database (31) to identify homologous genes in other phylogenetically related genomes, and we compared syntenic genomic regions for four representative genes which contain TE-derived DNA.

For example, the maize Zm-CACTA-181 TE is located in the 5-prime region of the gene Zm000021d052675, encoding for the sequence of the last seven gene exons. However, in the syntenic genomic region of *Brachypodium distachyon*, the orthologous gene misses all exons encoded in the TE sequence (**Figure 4A**). Similarly, the Zm-Pack-PIF-686 element has inserted into the Zm00001d036443 locus, encoding the gene’s second exon sequence. This insertion appears to be absent in the ortholog identified in *Setaria italica* (Si007332m.g, with ∼80% sequence identity), which lacks the exon located in the TE (**Figure 4B**). A Mariner Pack-TYPE element (Os-Mariner-4) contributes the fourth exon of the gene LOC_Os01g04110 in the rice genome. Also, in this case, the TE is not present in the *B. distachyon* ortholog (BRADI_2g02180v3), where the exon encoded within the TE is missing (**Figure 4C**). Finally, the Zm-Ac-1702 element has inserted into the maize gene Zm00001d025705, providing the splice donor site of the eighth exon of the gene, as well as 66 bp of its transcript (this fragment of coding DNA was too small to classify the element as Pack-TYPE with the parameters used in our analysis). In the syntenic DNA region of *Sorghum bicolor* (>80% sequence identity), the insertion of Zm-Ac-1702 is absent, and the orthologous gene’s exon terminates earlier (**Figure 4D**). In all these examples, the insertion of the Pack-TYPE element likely occurred after the phylogenetic separation of the related species, supporting the idea that all Pack-TYPE TEs can have exon shuffling activity associated with their mobilisation.

**Figure 4.**
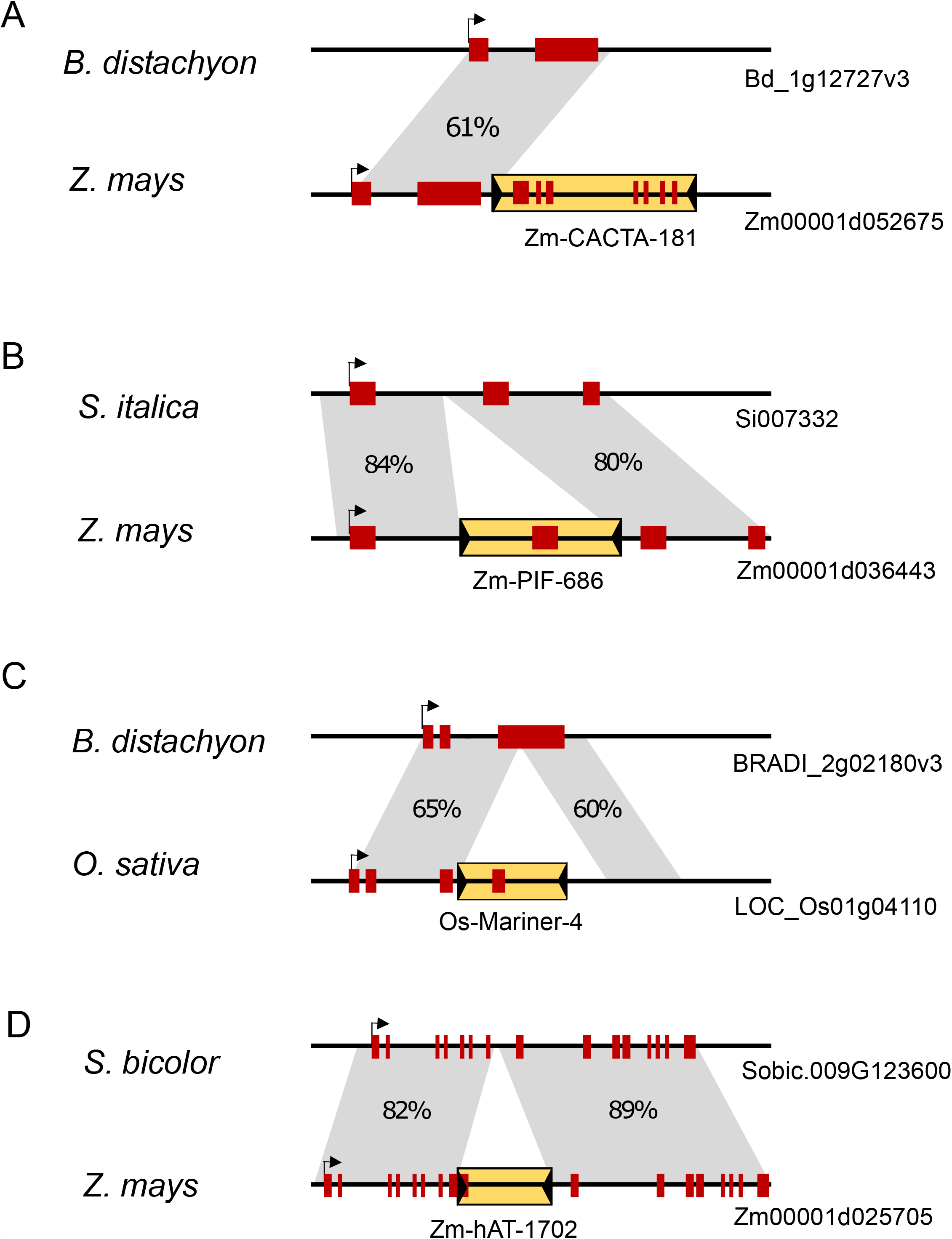
Pack-TYPE TEs contribute to maize gene evolution. **A** Scheme illustrating the maize Zm00001d052675 locus (with an insertion of a Pack-CACTA element) compared to the corresponding *B. distachyum* syntenic region. Sequence identity is displayed in the grey areas. The red blocks represent exons, and a black arrow indicates the start of transcription. The Pack-TYPE element is displayed as a yellow box with black triangles representing TIRs. **B** The Zm00001d036443 locus with the insertion of a Pack-PIF element, compared to a *S. italica* syntenic region. **C** The LOC_Os01g04110 locus with the insertion of a Pack-Mariner element, compared to a *B. distachyum* syntenic region. **D** The Zm00001d025705 locus with the insertion of a Pack-hAT element, compared to a *S. bicolor* syntenic region.

## Discussion

Several groups of TEs can mobilise and rearrange coding DNA in plant genomes (2), and there are several reported instances of gene transduplication events triggered by TEs (32). However, many of these cases are associated with complex and atypical transposition events, believed to occur rarely in nature, which have been selected as a consequence of their positive effects on gene expression. Only two TE superfamilies are reportedly able to systematically acquire coding DNA, including the Pack-MULEs described in rice (8, 33) and Helitron TEs in maize (34). Here, we provided evidence that not only MULEs but all main superfamilies of TIR-related TEs (i.e. CACTA, Mariner, PIF and hAT) can potentially generate Pack-TYPE elements in plants. The number of Pack-TYPE TEs we identified was variable in the maize and rice genomes and roughly correlated with the general abundance of TIR TE superfamilies annotated on each reference (25, 35). Generally, we observed a positive correlation between the total number of Pack-TEs detected in each group and their average length, which could be explained by the fact that longer elements are more likely to contain protein coding DNA.

Considering that most automatic annotation procedures for TIR TEs are homology-based (25, 36), these are not optimised to detect TEs with more variable DNA and limited similarity to known TE reference libraries, which is a common feature of Pack-TYPE TEs. Indeed, we found that even in the *A. thaliana* TAIR10 annotation, only 2% of Pack-CACTA elements were correctly identified as elements belonging to CACTA (ATENSPM) superfamily (**Figure 1**). Systematic studies of Pack-TYPE TEs have been reported only for Pack-MULE (9, 11, 33) as their relatively long TIR sequences facilitate their detection based on homology with existing MULE TEs (8, 10). Our approach uses a core TIR DNA sequence of 8 to 13 nucleotides as input; it relies on the conservation of TSDs and the presence of similar elements (clustering) as part of the detection process, allowing for the annotation of TE superfamilies with low TIR sequence conservation. In addition, contrary to most other TE annotation tools currently available, our detection pipeline is wholly embedded in an R/Bioconductor package (18) and can be easily performed using a single function, facilitating the detection and the study of Pack-TYPE transposons for any basic R user.

Besides Pack-TEs, we also annotated non-autonomous elements classified as non-Pack TEs; many of these belonged to the Mariner and PIF superfamilies in rice, and the PIF and hAT superfamilies in maize (**Figure 2**). Considering that our annotation analysis bases the discrimination between Pack-TYPE and non-Pack TEs on a BLAST search, it is possible that Pack-TYPE elements with particularly diverged sequences are not annotated as such simply because a significant BLAST hit could not be found.

Pack-TYPE TEs can potentially generate conflicts in the epigenetic regulation of the DNA portion with homology to genes because small RNAs produced to silence TEs can target similar sequences found in actively expressed genes (37–39). This phenomenon can lead to the fast pseudogenisation of less-constrained genes and rapid divergence of DNA sequences shared by functional genes and TEs (39), reducing the probability of significant blast hits. Therefore, BLAST approaches may only efficiently identify relatively recent acquisition events of protein coding DNA. Consistently, it has also been reported that the estimated age of Pack-MULE in rice tend to be young, with elements remaining recognisable only for a few million years (9).

On the other hand, the genomic location of the original neighbouring insertions primarily determines the DNA captured by Pack-TYPE TEs and may not include coding DNA (**Figure 3A**). While previous studies on Pack-MULEs in rice suggest the preferential insertion of these elements into genes (9, 10), it is also possible that at least a portion of the longer non-Pack TEs originated from capture events of intergenic chromosomal regions. In this case, these should be considered structurally similar to Pack-TYPE TEs. Nonetheless, considering that most TEs classified as non-Pack are less than 500 bp in length (**Figure 2B**), a large proportion are likely constituted by just the two TIRs. Elements with such features are known as Miniature Inverted TEs (MITEs) and are abundant in plant genomes, primarily associated with the PIF, Mariner, and hAT superfamilies (40–42).

Both MITEs and Pack-TYPEs are common among TIR TEs and seem to share similar structures and transposition mechanisms, likely competing for the same transposases. Nonetheless, their abundance in the genome varies depending on the TE superfamily and plant species considered. Therefore, considering the role of Pack-TYPE TEs in transduplication, the balance in the relative mobilisation of MITEs and Pack-TYPE TEs could directly affect plant genome plasticity and the speed of gene evolution.

## Data Availability

The entire set of annotated TEs are available from **Table S2**. These annotations were generated using packFinder v1.2.0, available as part of the Bioconductor project (https://doi.org/doi:10.18129/B9.bioc.packFinder). The code for the package may be found on GitHub (https://github.com/jackgisby/packFinder).

## Acknowledgements

Part of the computations described in this paper were performed using the University of Birmingham’s Compute and Storage for Life Sciences (CaStLeS) service.

## Funding

This work was partially funded by the Royal Society Research Grant [RGS\R1\201297].

## Conflict of interest

The authors declare no conflict of interest.

## Supplementary Table Legends

**Table S1**. CACTA-like elements identified in *Arabidopsis thaliana*.

**Table S2**. Elements identified with PackFinder in the *O. sativa* and *Z. mais* genomes.

**Table S3**. The number of non-overlapping maize and rice loci captured by each Pack-TE identified by BLAST.

**Table S4**. TEs used to produce the phylogenetic trees displayed in **Figure 3**.

**Table S5**. The number of Pack-TYPE TEs overlapping genes.

## Supplementary Figure Legends

**Figure S1.**
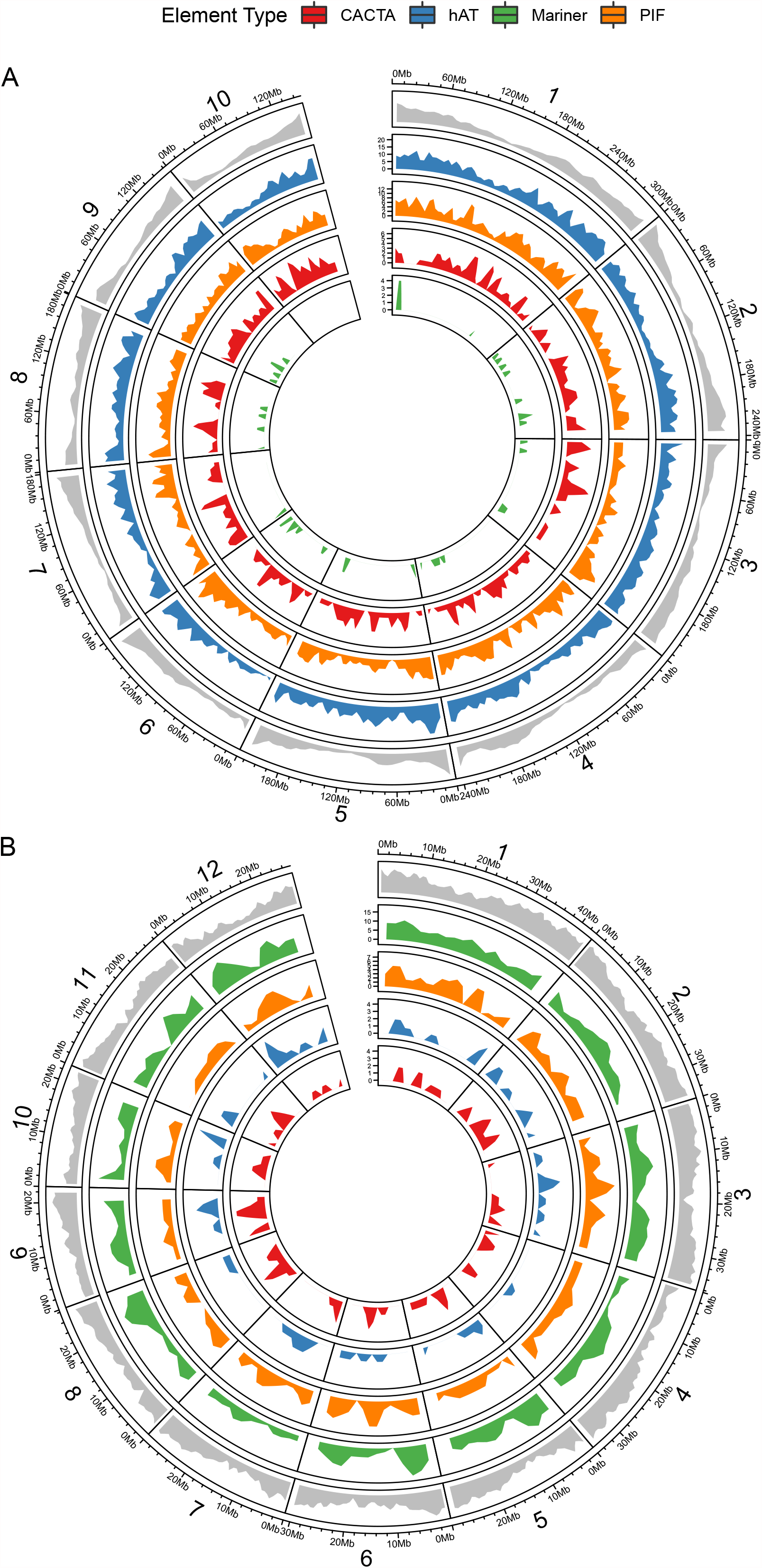
The whole-genome distribution of annotated TEs. Density plots, generated using Circos, for the distribution of the annotated TEs in *Zea mays* (**A**) and *Oryza sativa* (**B**). TE superfamilies are coloured and separated by track, ordered by their relative abundance in the genome. The distribution of genes (in grey) is plotted for comparison. The outer track displays genomic positions while the horizontal axis refers to TE frequency.

**Figure S2.**
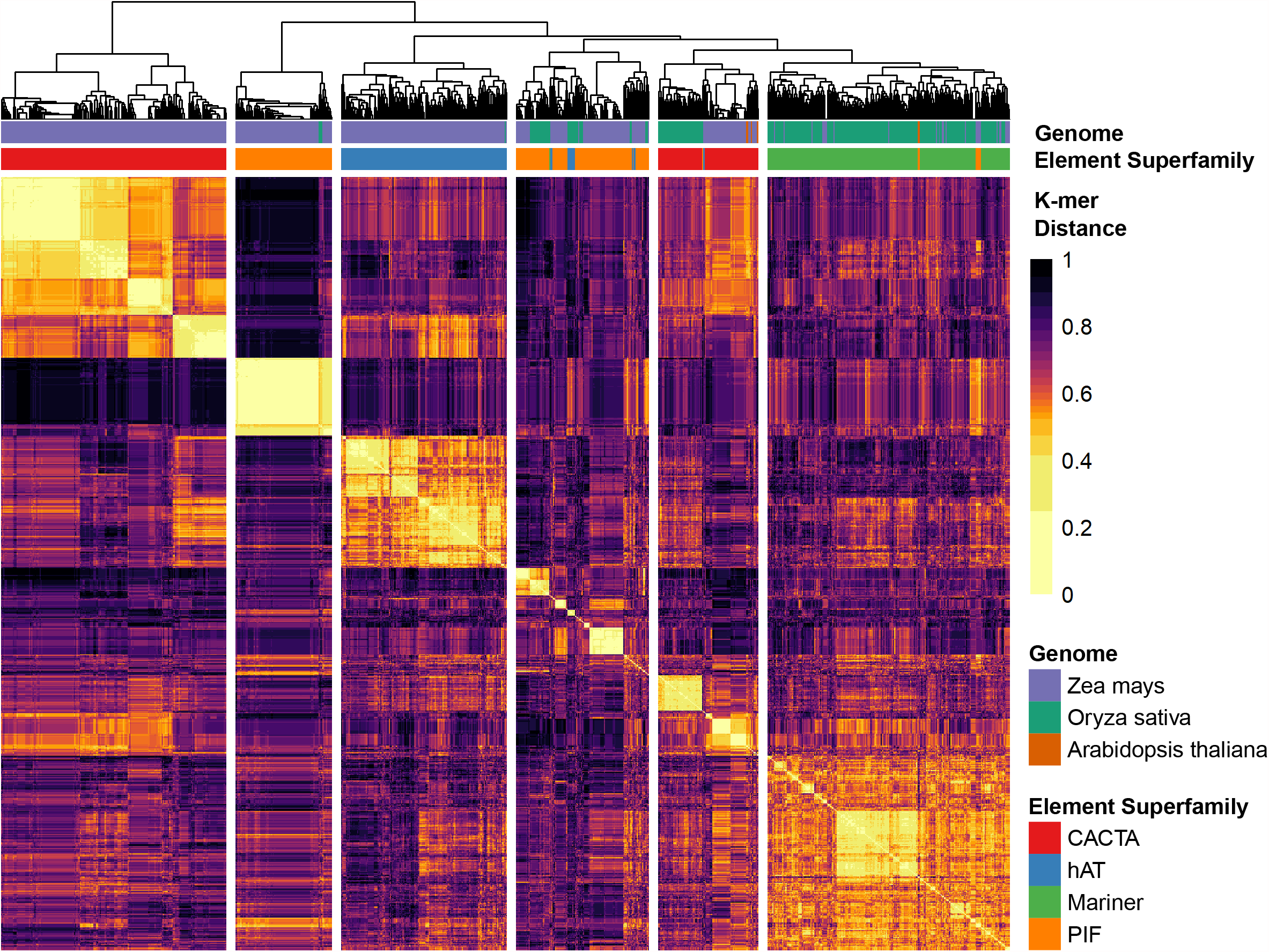
TIR relationships of annotated Pack-TYPE TEs. Visualisation of the symmetric distance matrix for the forward TIRs of all annotated Pack-TYPE TEs. Distance was calculated using the alignment-free kmer algorithm, and elements are ordered by hierarchical clustering (see **Methods**). We cut the dendrogram such that six groups were identified, separated by vertical whitespace.

**Figure S3.**
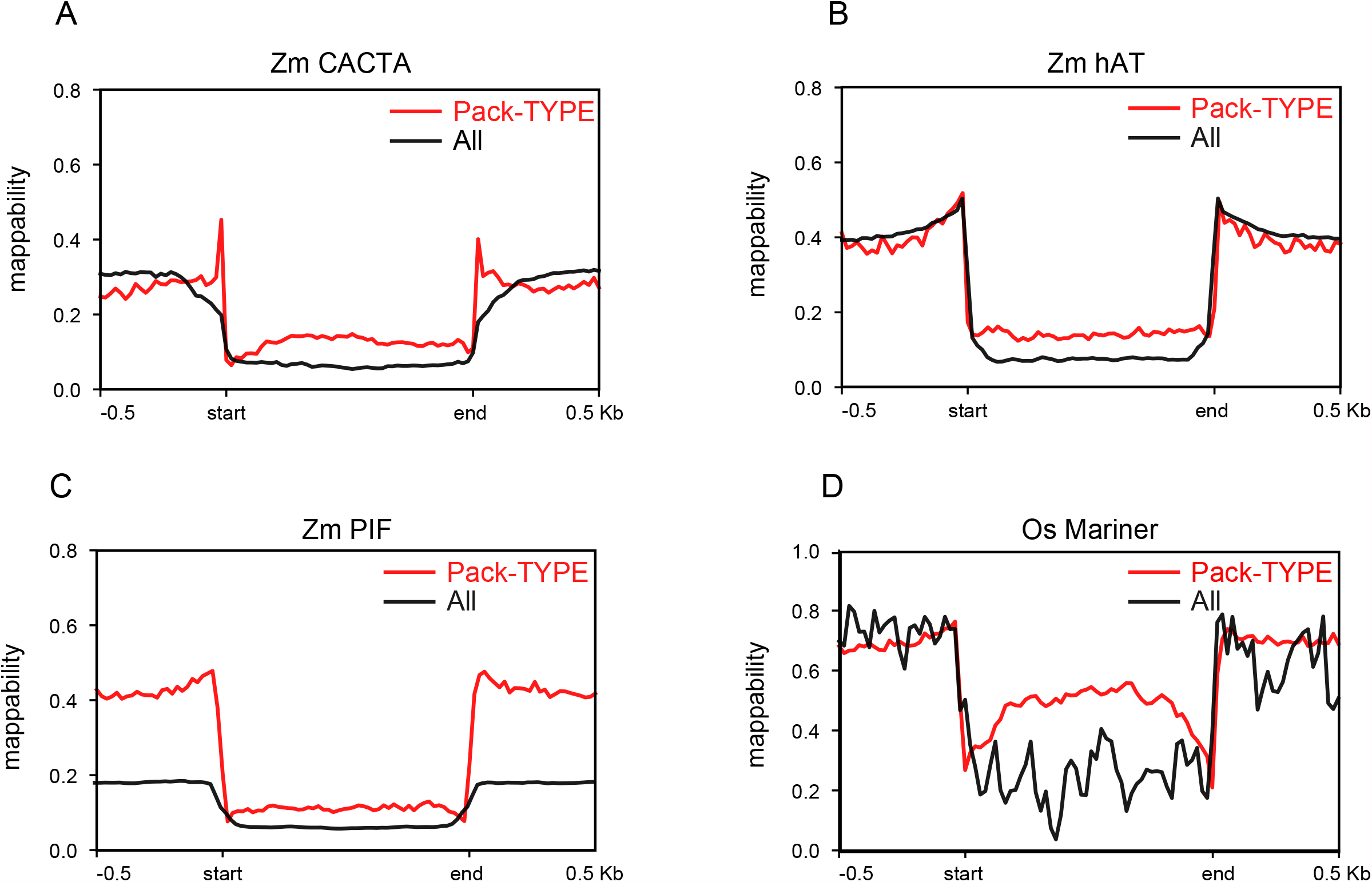
Repetitiveness of annotated Pack-TYPE-TEs. Distribution of averaged uniqueness of DNA sequence (mappability, see Methods), calculated for Pack-TYPE elements belonging to CACTA (**A**), hAT (**B**) and PIF (**C**) superfamilies in maize and for Pack-Mariner (**D**) in rice. The averaged mappability of all annotated elements belonging to each superfamily in the relevant genome (marked with “ All”) have been plotted for comparison.

**Figure S4.**
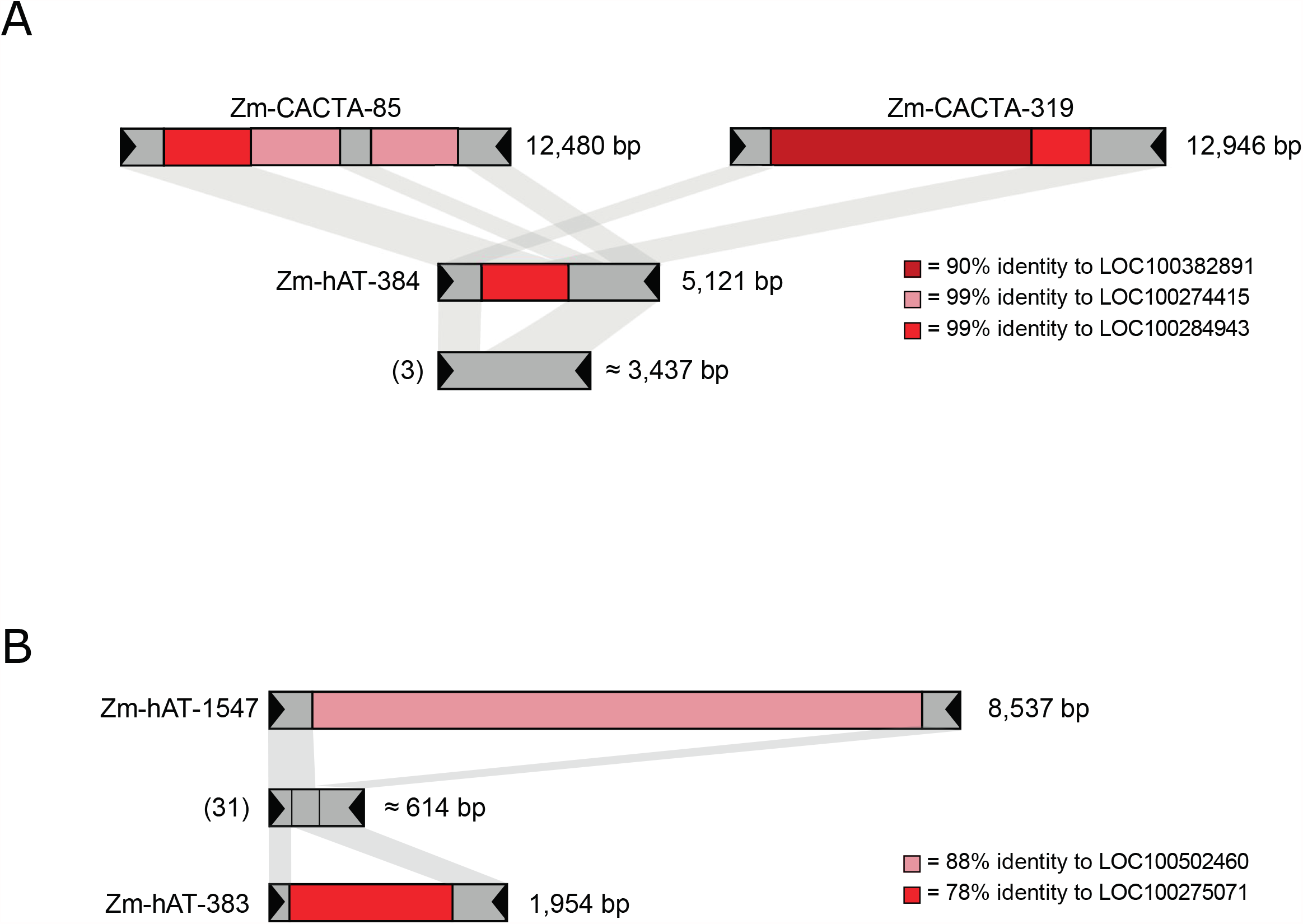
Examples of sequence structure of Pack-CACTA and Pack-hAT TEs. Each bar summarises the structure of a group (with the number of elements displayed in brackets) or single elements. Shaded grey marks regions with sequence homology. The size of elements is displayed on the right. Regions in different shades of red mark DNA sequences containing homology to coding genes. The black triangles mark the TIRs (15 nt). **A** Structure of the Zm-Pack-CACTA TE family #13. **B** Structure of the Zm-Pack-hAT TE family #14.

## Notes

### Competing Interest Statement

The authors have declared no competing interest.

